# Neural interactions in developing rhythmogenic spinal networks: Insights from computational modeling

**DOI:** 10.1101/2020.09.15.298281

**Authors:** Natalia A. Shevtsova, Ngoc T. Ha, Ilya A. Rybak, Kimberly J. Dougherty

**Author notes:** Correspondence: Ilya A. Rybak, Kimberly J. Dougherty.

## Abstract

The mechanisms involved in generation of rhythmic locomotor activity in the mammalian spinal cord remain poorly understood. These mechanisms supposedly rely on both intrinsic properties of constituting neurons and interactions between them. A subset of Shox2 neurons was found to contribute to generation of spinal locomotor activity, but the possible cellular basis for rhythmic bursting in these neurons remains unknown. Ha and Dougherty (2018) recently revealed the presence of bidirectional electrical coupling between Shox2 neurons in neonatal spinal cords, which can be critically involved in neuronal synchronization and generation of populational bursting. Gap junctional connections found between functionally-related Shox2 interneurons decrease with age, possibly being replaced by increasing interactions through chemical synapses. Here, we developed a computational model of a heterogeneous population of neurons sparsely connected by electrical or/and chemical synapses and investigated the dependence of frequency of populational bursting on the type and strength of neuronal interconnections. The model proposes a mechanistic explanation for emergence of a synchronized rhythmic activity in the neuronal population and provides insights into the mechanisms of the locomotor rhythm generation.

## 1 Introduction

The mammalian spinal cord contains neuronal circuits that can generate locomotor-like oscillations in the absence of supraspinal and afferent inputs. These circuits include rhythm-generating kernels capable of producing, maintaining and coordinating populational rhythmic activity. A rhythmogenic kernel on each side of the cord is thought to be comprised of mutually connected excitatory neurons with ipsilateral projections (Kjaerulff and Kiehn, 1996; Kiehn, 2006; Hägglund et al., 2010). Although no single genetically identified neuron type has been found to be solely responsible for rhythm generation in the spinal cord, the genetically identified Shox2 interneurons have been suggested to be involved in locomotor rhythmogenesis (Brownstone and Wilson, 2008; Dougherty et al., 2013). Later investigations provided evidence that a subpopulation of Shox2 neurons, specifically the Shox2 nonV2a neuron type, may belong to the rhythm-generating kernel in the neonatal rodent spinal cord and be involved in generation of locomotor rhythmic activity (Dougherty et al., 2013; Kiehn, 2016).

Although the existing data on the possible cellular basis for rhythmic bursting in the Shox2 neurons remains limited, there is evidence for the expression of several types of potentially rhythmogenic currents in these neurons, including the persistent inward current (Ha et al., 2019). Based on indirect data from multiple groups (Zhong et al., 2007; Tazerart et al., 2007, 2008; Ziskind-Conhaim et al., 2008; Brocard et al., 2010, 2013; Tong et al., 2012) and previous computational models (Rybak et al., 2006a,b; McCrea and Rybak, 2007; Sherwood et al., 2011; Zhong et al., 2012; Brocard et al., 2013; Rybak et al., 2013, 2015; Shevtsova et al., 2014; Danner et al., 2016, 2017, 2019; Shevtsova and Rybak, 2016; Ausborn et al., 2019), we have suggested that the generation of locomotor rhythmic activity in the spinal cord relies on the persistent sodium current (*I*_NaP_).

Regardless of the intrinsic bursting properties of single neurons involved in rhythmogenesis, mutual interactions between the neurons are critical to allow synchronization of neuronal activity leading to the generation of populational rhythmic activity. A recent study has shown that functionally related Shox2 interneurons in neonatal mice are locally connected with each other bidirectionally via electrical synapses or gap junctions (Ha and Dougherty, 2018). This bidirectional electrical coupling may represent a mechanism for neuronal synchronization and for populational oscillations in the neonatal spinal cord. Bidirectional gap junctional coupling within each population of genetically identified neurons, i.e. Shox2 nonV2a neurons, is found with high incidence in the neonatal spinal cord. With age, this electrical coupling decreases in both incidence and strength, which may occur in parallel (and possibly in connection) with an increase of the unilateral connections via excitatory chemical synapses (Ha and Dougherty, 2018). This change in the type, probability and strength of neuronal interactions may lead to specific changes in some characteristics of locomotor rhythm and pattern generated and the range of generated frequencies.

To theoretically investigate the potential roles of different (electrical and excitatory chemical) connections between Shox2 interneurons and their interaction with the *I*_NaP_ in generation of rhythmic populational activity, we developed a model of a population of neurons incorporating *I*_NaP_ in a subset of cells that are sparsely connected by electrical and/or excitatory chemical synapses and investigated the dependence of frequency of populational bursting on the connection type and strength, the maximal conductance and distribution of *I*_NaP_, and other neuronal characteristics. We demonstrated that the frequency of populational bursting activity depends on the relative expression of particular types of connections between neurons. The model proposes a mechanistic explanation for emergence of a synchronized rhythmic activity in a population of Shox2 neurons and provides insights into the mechanisms of the locomotor rhythm generation.

## 2 Results

The major focus of the present study was on the specific roles of, and possible cooperation between, neural interactions (through gap junction coupling and/or excitatory chemical synapses) and persistent (slowly inactivating) sodium current (*I*_NaP_) in neuronal synchronization and generation of populational rhythmic activity related to the locomotor-like oscillations in the mammalian spinal cord. We started by simulating a pair of neurons operating in different regimes and connected by bidirectional gap junctions or unidirectional excitatory chemical synapses in order to investigate the effect of the type and strength of connections and the presence/absence of *I*_NaP_ conductance on the activity of both neurons. Then, we considered a heterogeneous population of 100 neurons which incorporated a subpopulation of neurons containing *I*_NaP_ and hence could intrinsically generate rhythmic bursting depending on neuronal excitability. The remaining neurons were simple spiking neurons. The neurons had parameters randomly distributed over the population and were sparsely connected by electrical and excitatory chemical synapses. The probabilities of connections between neurons were set based on the data from Shox2 neurons (Dougherty et al., 2013; Ha and Dougherty, 2018). Our modeling study focused on neuronal interactions, synchronization of neuronal activity, and control of frequency of populational bursting depending on the relative expression of different types of neuron connectivity and connection strength.

### 2.1 Gap junctional coupling in two-cell model

We first simulated a pair of interneurons (In1 and In2) modelled in the Hodgkin-Huxley style and coupled bidirectionally by gap junctions with conductance *g*_Gap._ Each neuron represented either a conditionally bursting neuron with *I*_NaP_ or a simple tonically spiking neuron without *I*_NaP_. The equations describing neuron dynamics and neuron parameters are specified in **Materials and Methods**.

The baseline operating regime of each neuron (prior to coupling) was dependent on its resting membrane potential, which in turn was defined by the value of the leakage reversal potential, *E*_L_ (see **Table 1**). With an increase in *E*_L,_ the isolated conditionally bursting neuron changed its operating regime from silence to bursting and then to tonic spiking. The isolated “simple” neuron without *I*_NaP_ could switch from silence to tonic spiking when *E*_L_ increased. The strength of gap junctional coupling between the neurons (*g*_Gap_) was varied. The time course of the neuron membrane potentials for various baseline operating regimes and the strength of coupling is illustrated in **Figure 1**. The frequencies of bursting, as well as neuron resting potentials at various values of *g*_Gap_ are shown in **Table 1**.

**Table 1.** Effect of electrical coupling on neuron activity in two-cell model

| Neuron | $I_{NaP}$ | $E_L$ | Regime | Resting potential, mV | | | | Frequency, Hz | | | |
| --- | --- | --- | --- | --- | --- | --- | --- | --- | --- | --- | --- |
| | | | | Connection strength ($g_{Gap,nS}$) | | | | Connection strength ($g_{Gap,nS}$) | | | |
|  |  |  |  | 0 | 0.05 | 0.1 | 0.2 | 0 | 0.05 | 0.1 | 0.2 |
| In1 | + | -69 | B | -66 | -66 | -66.1 | -66.4 | 0.2 | 0.2 | **0.16** | **0.14** |
| In2 | + | -72.2 | S | -71 | -70.5 | -70.2 | -70 | N/A | 0.08 |  |  |
| In1 | + | -69 | B | -66 | -66.4 | -66.4 | -66.4 | 0.2 | 0.2 | **0.35** | **0.33** |
| In2 | + | -65.5 | B | -62.7 | -62.8 | -62.9 | -62.9 | 0.43 | 0.36 |  |  |
| In1 | + | -69 | B | -66 | -65.8 | -67 | -66.5 | 0.2 | 0.2 | **0.39** | **0.36** |
| In2 | + | -64.9 | T | -55.4 | -61.8 | -61.9 | -62.7 | N/A | 0.44 |  |  |
| In1 | + | -71.4 | S | -69.3 | -67 | -66.5 | -66 | N/A | **0.05** | **0.1** | **0.13** |
| In2 | - | -59.6 | S | -57 | -58.5 | -59.1 | -59.9 | N/A |  |  |  |
| In1 | + | -69 | B | -66 | -65.8 | -65.6 | -65.2 | 0.2 | **0.22** | **0.23** | **0.25** |
| In2 | - | -59.6 | S | -57 | -58.3 | -58.7 | -59.8 | N/A |  |  |  |
| In1 | + | -69 | B | -66 | -65.8 | -65.6 | -65 | 0.2 | **0.22** | **0.24** | **0.26** |
| In2 | - | -59.3 | T | -56.1 | -57.7 | -58.5 | -58.3 | N/A |  |  |  |

“+” – neuron incorporating the persistent sodium current ( $I_{NaP}$ ); “-” – neuron non-incorporating $I_{NaP}$ ; “B” – neuron generating bursting activity; “S” – neuron in a silent mode; “T” – neuron generating tonic activity; frequency of synchronized activity of two neurons is shown by bold.

**Figure 1.**
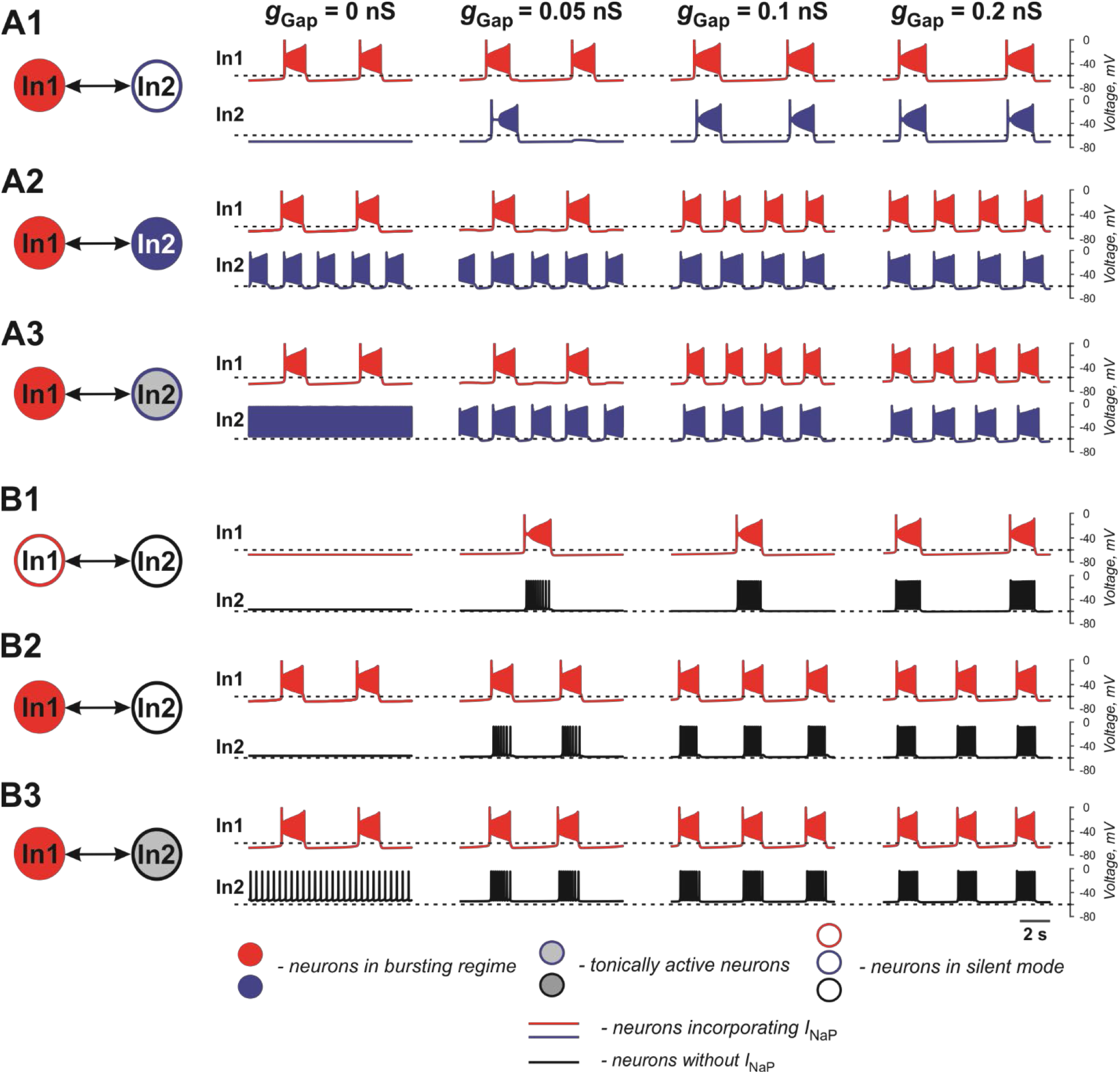
Effects of electrical coupling on activity of two neurons (In1 and In2). **(A1-A3)** Both neurons incorporate the persistent sodium current (*I*_NaP_). In1 is in a bursting mode and In2 is silent if uncoupled **(A1)**, operates in a bursing regime **(A2)**, or is tonically active **(A3). (B1-B3)** Only In1 incorporates *I*_NaP_ and, if uncoupled, is silent **(B1)** or operates in a busting regime (**B2** and **B3**). In2, if uncoupled, is either silent (**B1** and **B2**) or tonically active **(B3)**. The strength of electrical coupling increases from left to right.

In our simulations shown in **Figures 1A1-A3**, both neurons (In1 and In2) incorporate *I*_NaP_. In all three panels, In1, if uncoupled, operates in a bursting mode, whereas In2 is silent (at low level of excitation, panel A1), exhibits bursting (at intermediate level of excitation, panel A2), or is tonic spiking (at a highest level of excitation, panel A3). The effect of gap junctional coupling depends on the difference between the membrane potentials of coupled neurons and the strength of coupling, and hence can be depolarizing for one neuron and hyperpolarizing for the other one. In all three cases, gap junctional coupling leads to synchronization of neuronal bursting from the burst ratio M:1, M>1 at a lower *g*_Gap_ (0.375 < *g*_Gap_ < 0.66 nS) to the ratio of 1:1 at higher *g*_Gap_ (*g*_Gap_ ≥ 0.66 nS) values. Note that coupling may reduce bursting frequency relative to the uncoupled case (see panel **A1**) or increase it (panels **A2** and **A3**, see also **Table 1**) depending on the relative values of the resting membrane potentials of coupled neurons. This is seen in **Figure 1A1** when In1 depolarizes In2 and induces bursting in this neuron with common bursting frequency at higher *g*_Gap_. At the same time, In2 hyperpolarizes In1 resulting in a general decrease of common bursting frequency relative to the uncoupled case with increased *g*_Gap_ (see **Table 1**). In **Figures 1A2,A3**, we see the opposite situation. In these cases, both neurons operate in bursting regimes. However, the level of excitation of In1 is lower than that of In2, and prior to coupling, In2 generates bursts with a higher frequency than In1 in **Figure 1A2** and is tonically active in **Figure 1A3**. Electrical coupling of these neurons leads to depolarization of In1 and hyperpolarization of In2. As a result, burst frequency of In1 increases while burst frequency of In2 decreases. In **Figure 1A3**, at lower *g*_Gap_ In2 switches from tonic to bursting mode, when coupled, with the burst ratio 1:2. Similar to the simulation shown in **Figure 1A1**, at higher *g*_Gap_ the two-cell model shows a synchronized bursting activity; however, the resulting frequency is higher than the initial In1 frequency (see **Table 1**). Hence gap junctional coupling between neurons with *I*_NaP_ leads to synchronized rhythmic bursting with bursting frequency dependent on neuronal excitability and the strength of coupling.

In simulations shown in **Figures 1B1-B3**, only one of the two electrically coupled neurons (In1) has *I*_NaP_. Prior the coupling, this neuron can be silent (panel **B1**) or operate in a bursting regime (panels **B2** and **B3**), whereas the other, “simple” neuron (In2), depending on the level of its excitation, can be silent (panels **B1** and **B2**) or exhibit tonic spiking (panel **B3**). In all these simulations, the In2 neuron has higher resting potential if uncoupled and depolarizes In1 if the two neurons are connected by gap junctions (see **Table 1**). This results in the emergence of bursting activity (as in panel **B1**) or in an increase of the bursting frequency with increasing *g*_Gap_ (panels **B2** and **B3**). In all three cases, In1 hyperpolarizes In2 during interburst intervals but induces high-frequency spiking in In2 during bursts, resulting in synchronized activity of the two-cell model replicating bursting activity of In1.

The two neurons in **Figure 2** are similar to those shown in **Figure 1B1** and only one of the two electrically coupled neurons (In1) has *I*_NaP_. Here, instead of varying the strength of gap junctional coupling, it was set at 0.1 nS and the effects of varying the strength of *I*_NaP_ are illustrated. At 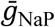 below the level shown in **Figure 2**, both neurons remain silent. However, with increasing 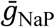, bursting emerges and increases in frequency. The results demonstrate that the presence of *I*_NaP_ in one neuron can initiate synchronized bursting in electrically coupled neurons even if both neurons when uncoupled are silent.

**Figure 2.**
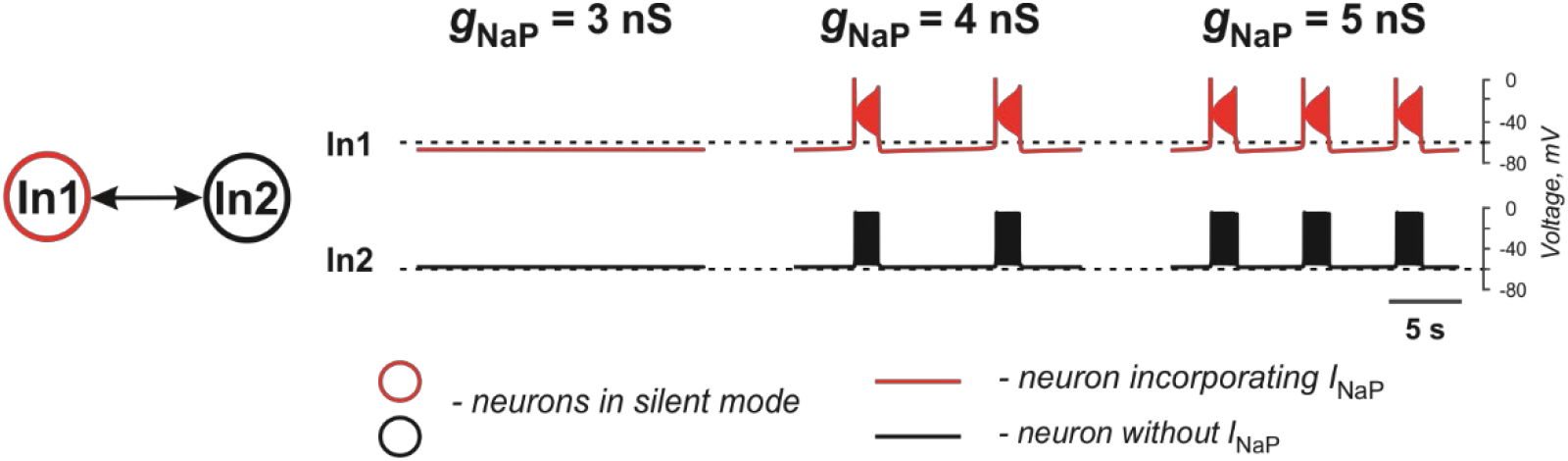
Effects of increasing *I*_NaP_ conductance (*g*_NaP_) on activity of two neurons mutually connected via gap junction. In these simulations, *E*_L1_ = -71.4 mV, *E*_L2_ = -58.4 mV, and the strength of the electrical connection (*g*_Gap_) is 0.1 nS.

Overall, gap junctional coupling leads to equilibration of membrane potentials between the coupled neurons, regardless of the presence/absence of *I*_NaP_. Depending on the direction of change (depolarization or hyperpolarization), this coupling will either increase or decrease bursting frequency of coupled neurons with *I*_NaP_ and bring neurons without *I*_NaP_ from silent or tonic to bursting mode in synchrony with neurons with *I*_NaP_. In turn, *I*_NaP_ in some neurons can amplify the effect of electrical coupling of these neurons with other neurons. Finally, both the presence of *I*_NaP_ in some neurons and electrical coupling between neurons promotes neuronal synchronization and both are critical for initiation and support of populational bursting.

### 2.2 Excitatory chemical synaptic connections in two-cell model

In the case of excitatory connections by chemical synapses, we considered only unidirectional connections because reported chemical synapses between spinal interneurons have been unidirectional (Dougherty et al., 2013; Ha and Dougherty, 2018; Chopek et al. 2018). **Figure 3** and **Table 2** illustrate effects of excitatory chemical synaptic connections in the two-cell model when both neurons (source neuron In1 and target neuron In2) are bursting neurons incorporating *I*_NaP_ (panels **A1-A3**), or when *I*_NaP_ is incorporated in only In1 (panels **B1** and **B2**) or only In2 neuron (panels **C1** and **C2**).

**Table 2.** Effect of chemical coupling on neuron activity in two-cell model

| Neuron | $I_{NaP}$ | $E_L$ | Regime | Resting potential, mV | | | | Frequency, Hz | | | |
| --- | --- | --- | --- | --- | --- | --- | --- | --- | --- | --- | --- |
| | | | | Connection strength ($w_{Syn}$) | | | | Connection strength ($w_{Syn}$) | | | |
|  |  |  |  | 0 | 0.5 | 1 | 2 | 0 | 0.5 | 1 | 2 |
| In1 | + | -65.5 | B | 62.7 |  |  |  | 0.43 |  |  |  |
| In2 | + | -71.4 | S | -69.3 | -70.5 | -63.1 |  | N/A | 0.2 | **0.43** |  |
| In2 | + | -69 | B | -66 | -68.3 |  |  | 0.2 | **0.43** |  |  |
| In2 | + | -64.9 | T | -55.4 | -61.7 |  |  | N/A | **0.43** |  |  |
| In2 | - | -59.6 | S | -57 |  |  |  | N/A | **0.43** |  |  |
| In2 | - | -59.3 | T | -56.1 |  |  |  | N/A | **0.43** |  |  |
| In1 | - | -58.4 | T | -55.2 |  |  |  | N/A |  |  |  |
| In2 | + | -72.2 | S | -71 | -70.5 | -70 | -68.5 | N/A | N/A | N/A | 0.12 |
| In2 | + | -69 | B | -66 | -65.6 | -65.1 | -64.5 | 0.2 | 0.22 | 0.25 | 0.33 |

Notation is the same as in Table 1.

**Figure 3.**
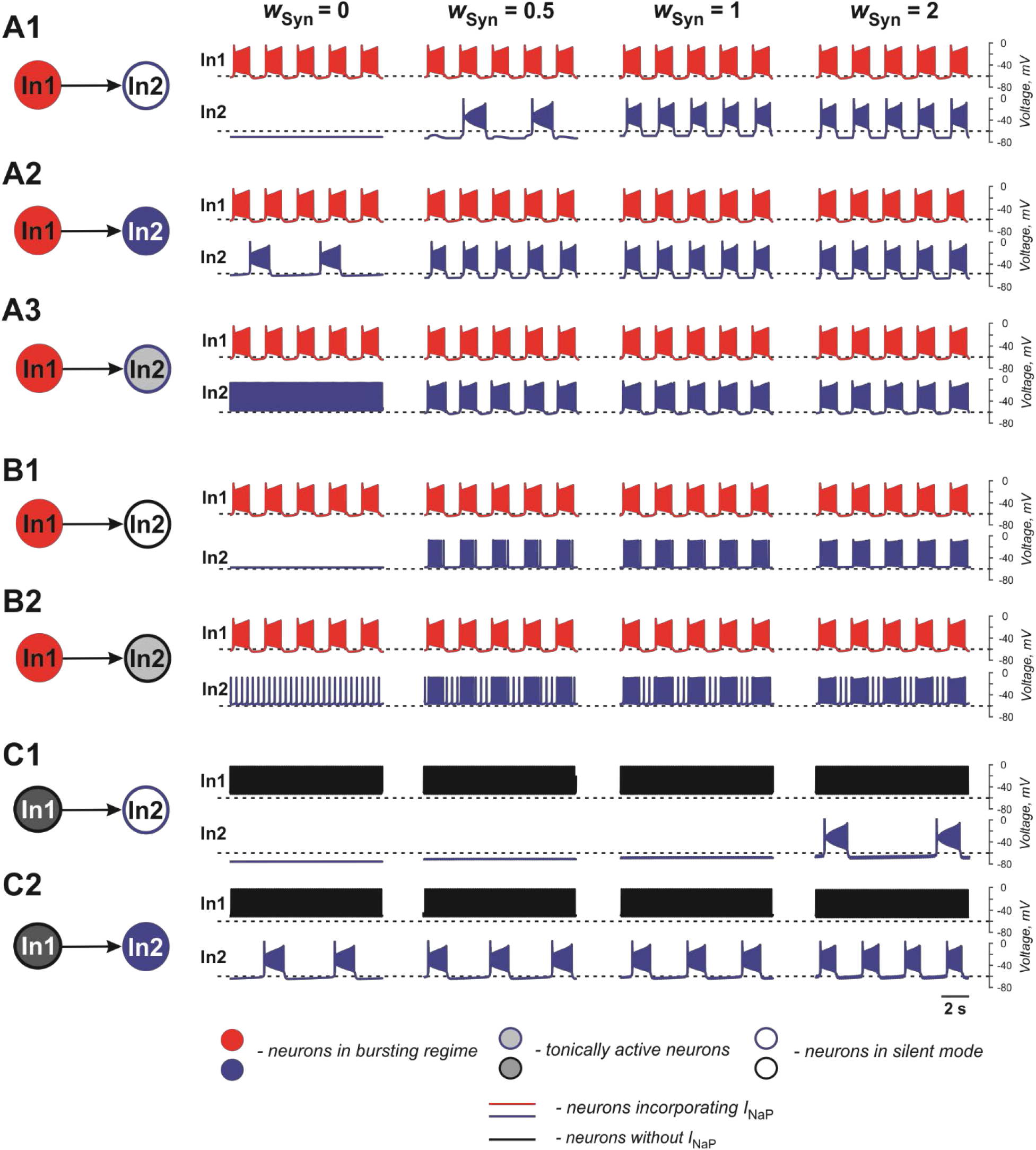
Effects of the excitatory chemical coupling on activity of two neurons (In1 and In2). **(A1-A3)** Both neurons incorporate the persistent sodium current (*I*_*NaP*_). In1 is in a bursting mode and In2, if uncoupled, is silent **(A1)**, or operates in a bursing regime **(A2)**, or is tonically active **(A3). (B1,B2)** Only In1 incorporates *I*_NaP_ and operates in a busting regime and In2, if uncoupled, is either silent **(B1)** or tonically active **(B2). (C1,C2)** In1 does not incorporate *I*_NaP_ and and is tonically active. In2 incorporates *I*_NaP_ and if uncoupled, is either silent **(C1)** or operates in a bursting regime (**C2**). The weight of synaptic connection increases from left to right.

In **Figures 3A1-A3**, the target neuron (In2) that incorporates *I*_NaP_ operates in various regimes without connection: silent (panel **A1**), bursting (panel **A2**), and tonic spiking (panel **A3**). The synaptic input from In1 to In2 leads to bursting activity in In2 with a burst ratio of 2:1 at a low *w*_Syn_ (0.5) or synchronized bursting of both neurons (at higher *w*_Syn_ values in panel **A1** and in panels **A2** and **A3**, see also **Table 2**).

In **Figures 3B1,B2**, the target neuron In2 does not have *I*_NaP_ and when uncoupled is either silent (panel **B1**) or generates tonic spiking (panel **B2**). In these cases, when In2 receives rhythmic excitatory synaptic input from the source neuron In1, it follows In1 activity and generates bursts synchronously with the source. In **Figure 2B2**, the bursting activity in In2 occurs on the background of baseline spiking activity.

In **Figures 3C1,C2**, the source neuron In1 is a “simple” neuron without *I*_NaP_ that generates a sustained spiking activity providing constant excitatory synaptic drive to the target In2 neuron with *I*_NaP_. This drive depolarizes In2 and this depolarization increases with the strength of synaptic input (*w*_Syn)_. Therefore, depending on the baseline regime of the target In2 neuron, this input results either in the emergence of bursting activity (panel **C1**) or the increasing of bursting frequency (panel **C2**, see also **Table 2**) in the target In2 neuron.

The above simulations have shown that chemical connections in the two-cell model promote neuronal synchronization and have variable effects on the resultant common bursting frequency.

### 2.3 Populational models

To study the synchronization and frequency control in a heterogeneous population of neurons connected by electrical and/or excitatory chemical synapses, we developed a model of a population of 100 neurons consisting of neurons with the intrinsic bursting properties (based on the incorporated *I*_NaP_) and “simple” neurons without such properties (with base ratio of 40/60%). All neurons were described in the Hodgkin-Huxley style as in the two-cell models described above. The neurons in the population were sparsely connected by electrical and/or excitatory chemical synapses with probability of *p*_Gap_ = 0.3 and *p*_Syn_ = 0.1 (Dougherty et al., 2013; Ha and Dougherty, 2018). Following Butera et al. (1999b), we chose the leakage reversal potential, *E*_L_, the leakage conductance, *g*_Leak_, and the maximum conductance of the persistent sodium current, 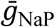, to be randomly distributed over population to provide natural heterogeneity between single neuron states. These parameters are the ones that define the basic level of neuronal excitation and their ability to intrinsically generate bursting activity (Butera et al. 1999a,b; Del Negro et al., 2002; Koizumi and Smith, 2008). *E*_L_, 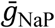 and *g*_Leak_ for neurons were distributed normally with a priory assigned means and standard deviations (see **Materials and Methods**). The major focus of our studies was on the generation of populational activity and frequency control of populational bursting for different types of connections and depending on the relative expression of different types of neuron connectivity and connection strength.

#### 2.3.1 Performance of populational model with electrical coupling

To investigate the role of gap junction in populational bursting, we simulated a population of 100 neurons with bidirectional electrical coupling with coupling probability of 0.3. **Figure 4** shows the model performance for different values of coupling (*g*_Gap_) in the population. In each panel, the upper row shows the averaged populational activity, the middle row shows raster plots of spikes elicited by all neurons in the population, and the bottom row shows traces of membrane potentials of several sample neurons with different excitabilities. Neurons with *I*_NaP_ are shown by red and neurons without *I*_NaP_ are shown by black. In **Figure 4A**, neurons in the population are uncoupled (*g*_Gap_ = 0). Because of the random distribution of neuronal parameters, neurons operate in different basic regimes: some neurons with *I*_NaP_ are silent, some generate bursting activity with different burst frequencies, and some are tonically active. Similarly, some neurons without *I*_NaP_ are silent and some demonstrate tonic spiking activity with different frequencies. The integrated neuron activity is sustained (upper row in **Figure 4A**). With increased electrical coupling (*g*_Gap_ = 0.033 nS, **Figure 4B**), most of neurons incorporating *I*_NaP_ synchronize their bursting activity and some neurons without *I*_NaP_ become involved in populational bursting. Further increase of *g*_Gap_ (*g*_Gap_ = 0.066 nS, **Figure 4C**) results in synchronization of bursting activity of all neurons incorporating *I*_NaP_ and involvement of the majority of neurons without *I*_NaP_ in populational bursting, which increases frequency and amplitude of populational bursting activity (see upper row in **Figure 4C**). However, after all neurons are involved in populational bursting, further increase of *g*_GaP_ does not lead to increasing frequency of oscillations. Instead, burst duration starts to increase and interburst intervals to decrease and eventually all neurons switch to sustained tonic activity (not shown).

**Figure 4.**
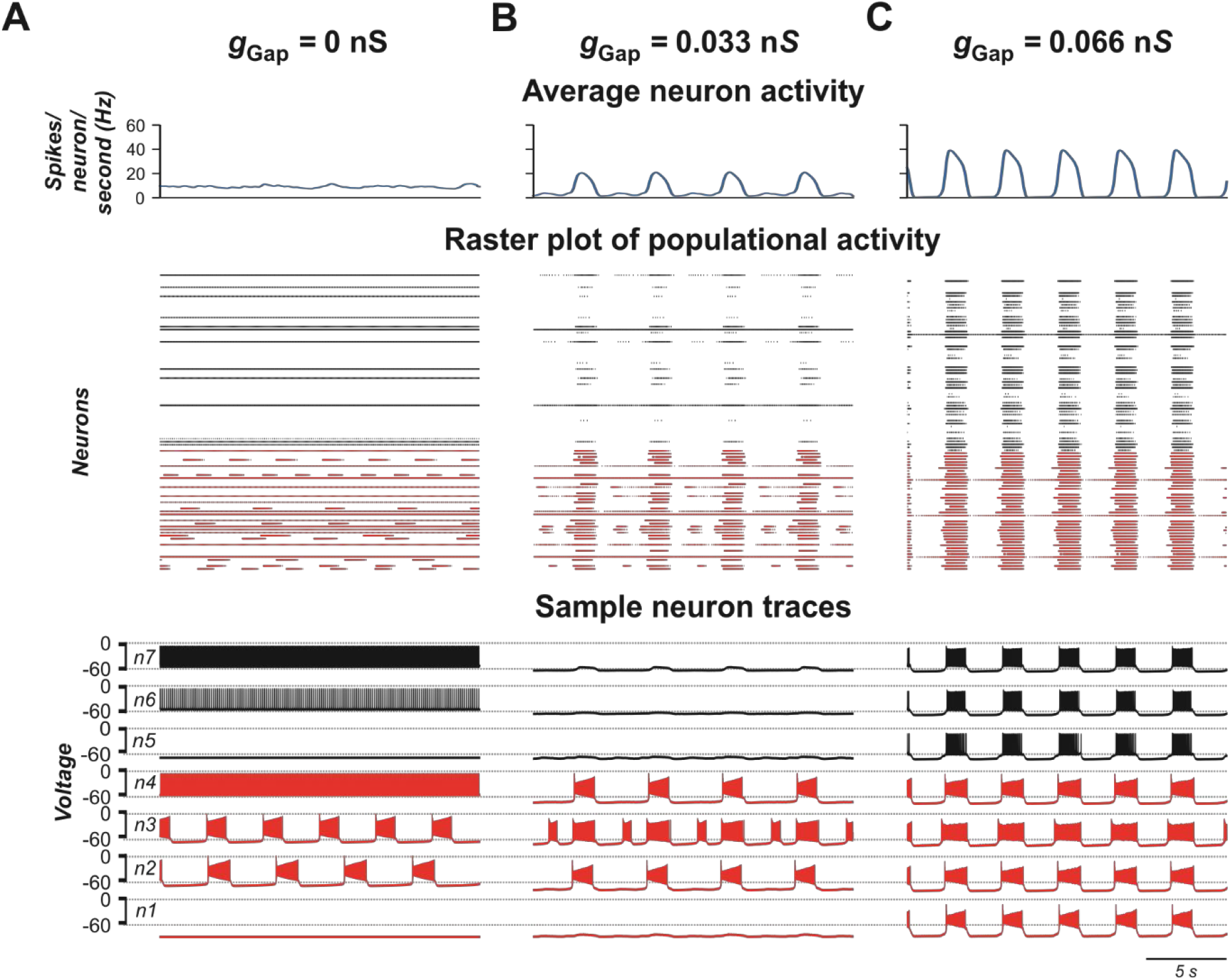
Model performance at different values of electrical connection strength in the population (*g*_Gap_). *g*_Gap_= 0 in **A**, 0.033 in **B**, and 0.066 in **C**. The probability of connection is 0.3. The upper row in each panel shows populational activity as an averaged histogram of neuron spiking [spikes/(*N* × s), where *N* = 100 is the number of neurons in population; bin = 100 ms]. The middle rows show raster plots of spikes elicited by neurons in the population. The lower rows show traces of the membrane potential of sample neurons. 40% of neurons in the population incorporate *I*_NaP_ and are shown by red and neurons without *I*_NaP_ are shown by black.

Interestingly, the frequency and amplitude of populational bursting depends on expression of *I*_NaP_ in neurons of the population. **Figure 5** shows the dependence of the frequency and amplitude of populational bursting on the average value of the *I*_NaP_ conductance 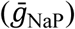 for three different values of *N*_NaP,_ defining the number of neurons with *I*_NaP_ in the population. As *N*_NaP_ increases, the populational bursting starts at lower values of 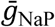. The burst frequency increases with increasing values of both 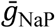 and *N*_NaP_ (panel **A**), and amplitude of populational bursts is higher at greater *N*_NaP_ values but depends on 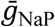 only at lower 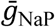 values (panel **B**).

**Figure 5.**
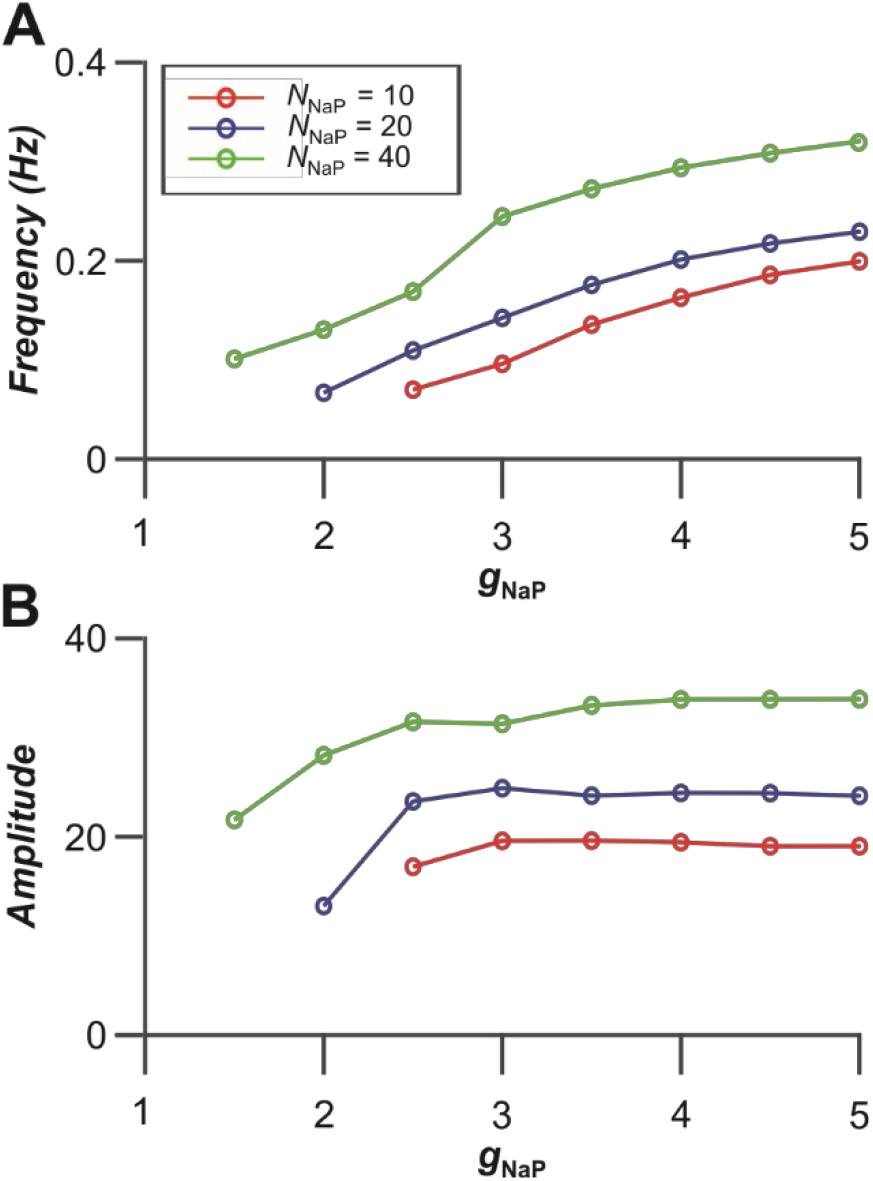
Dependence of bursting frequency **(A)** and amplitude **(B)** on the average value of the *I*_NaP_ conductance (*g*_NaP_) for three values of *N*_NaP_, the number of neurons incorporating *I*_NaP_ in the population. In these simulations, *g*_Gap_ = 0.06 nS.

#### 2.3.2 Gap junctional coupling and locomotor-like activity in the spinal cord

Previous experimental studies have shown that blocking gap junctions with carbenoxolone reduces the frequency of ventral root bursts of locomotor-like activity in the isolated rodent cord evoked by the mixture of NMDA and 5-HT (Ha and Dougherty, 2018). **Figures 6A1,A2** show an example of ventral root recordings during drug-evoked locomotion prior to and 30 min after carbenoxolone application begins. Burst frequency and amplitude are quantified in **Figures 6B1,B2** demonstrating that blocking gap junctions in these preparations reduces the frequency of locomotor oscillations and, in most cases, the amplitude of integrated locomotor activity. To simulate the effect of blocking electrical synapses, we reduced the strength of gap junctional coupling between neurons by 50% in our model. **Figures 6C1,C2** demonstrate the average populational activity for a sample simulation when *g*_Gap_ was reduced by 50%. To generally estimate the effect of this reduction on frequency and amplitude of synchronized bursting, we ran several simulations for populations with different randomization of neuron parameters. As can be seen in **Figures 6D1,D2**, regardless of parameter distribution, our model exhibits changes in the frequency and amplitude of oscillations with a decrease of *g*_Gap_ similar to experimental results shown in **Figures 6A1,A2,B1,B2**.

**Figure 6.**
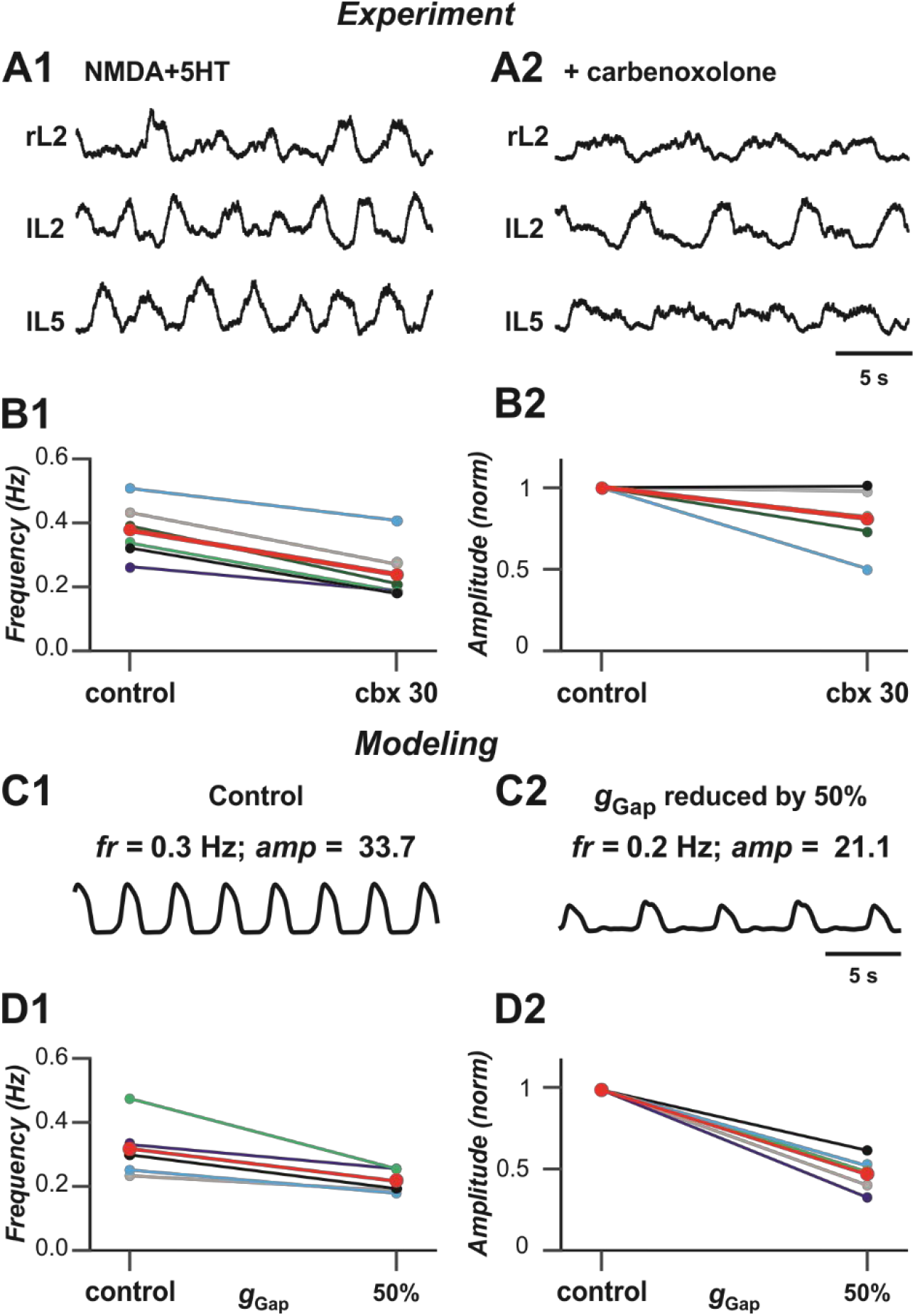
Blocking gap junctions decreases locomotor frequency. **(A1,A2)** Extracellular recordings from ventral roots at lumbar level 2 (L2)-flexor dominant root- and level 5 (L5)-extensor dominant root-on the right (r) and left (l) sides of the spinal cord after application of NMDA (7 mM) and serotonin (8 mM) in control case and after addition of carbenoxolone (100 μM). (**B1,B2)** Effect of carbenoxolone application on frequency **(B1)** and normalized amplitude **(B2)** of locomotor activity for 6 spinal cord preparations. Each circle represents the mean value of estimated parameter for one recording. Thinner lines connecting two circles represent parameter change in one cord. Thick red lines represent the means. The black lines in **B1** and **B2** correspond to the experimental recordings shown in **A1** and **A2. (C1,C2)** Effect of decreased strength of electrical coupling (*g*_Gap_) on populational activity. In **C2**, *g*_Gap_ is reduced by 50%. (**D1,D2)** Effect of *g*_Gap_ reduction on frequency **(D1)** and normalized amplitude **(D2)** of locomotor activity for 5 simulations with different randomizations of model parameters (simulation time 60 s). Each circle represents the average value of estimated parameter for one simulation. Light colored lines connecting two circles represent parameter change in simulation. Thick red lines represent the means. The black lines in **D1** and **D2** correspond to the simulations shown in **C1** and **C2**.

#### 2.3.3 Performance of populational model with chemical coupling

In the next stage of this study, we considered the same population of 100 neurons, but replaced the bidirectional gap junction connections within the population with unilateral excitatory chemical connections with probability of *p*_Syn_ = 0.1 (Dougherty et al., 2013). **Figure 7** shows the model performance for different weights of excitatory synaptic connections (*w*_Syn_) in the population. As can be seen in these simulations, enhancing *w*_Syn_ increases the number of tonically active neurons both among neurons with and without *I*_NaP_ (**Figures 7A,B**). Simultaneously, the bursting neurons start to consolidate in clusters including synchronously active neurons with *I*_NaP_ and partially involving neurons without *I*_NaP_ in bursting activity. Further increase of *w*_Syn_ increases frequency of bursting in neurons with *I*_NaP_ and synchronizes all neurons, resulting in populational synchronous bursting of high frequency and amplitude. In contrast to the population of electrically coupled neurons, in the population with chemical coupling, an increase in *w*_Syn_ is accompanied by a simultaneous increase of the background tonic activity.

**Figure 7.**
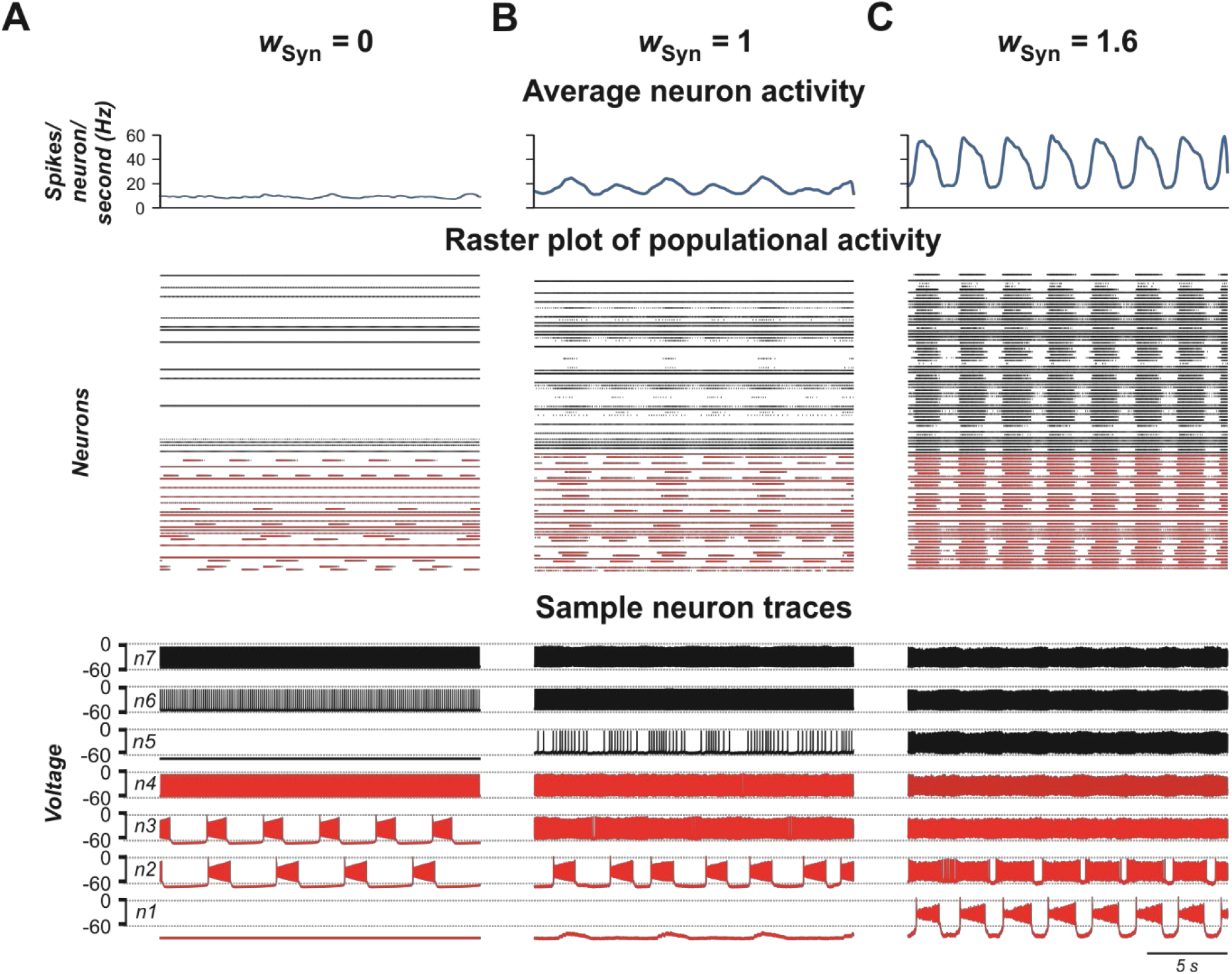
Model performance at different values of excitatory chemical connection strength in the population (*w*_Syn_). The probability of connection is 0.1. Arrangement and notation are the same as in **Figure 4**.

#### 2.3.4 Contribution of neuronal coupling by electrical and chemical synapses to populational activity

Experimental studies have shown that electrical coupling between Shox2 neurons is age-dependent. It is prevalent between functionally related Shox2 neurons in neonatal spinal cords and decreases in incidence and strength with age (Ha and Dougherty, 2018). To investigate how the activity of the population changes when both electrical and excitatory chemical connections are varied, we simulated a population in which both electrical and excitatory chemical synapses are present and sparsely distributed among neurons with probabilities of *g*_Gap_ = 0.3 and *p*_Syn_ = 0.1, and introduced two variables *G* and *W*, characterizing in average the total network electrical or chemical synaptic input to each neuron in the population, respectively (see **Materials and Methods**).

**Figures 8A,B** show 2D diagrams of color-coded frequency and amplitude of populational bursts in [*G, W*] plane. As seen in **Figure 8A**, if *W* is fixed and *G* is increasing, for most values of *W*, frequency of populational bursting does not increase smoothly but there are distinct areas in which the frequency does not change much and transition points when frequency increases abruptly. For example, at lower values of *W* (*W* < 10) there are one or two distinct transitions from the dark blue to light blue to cyan or the dark blue to cyan color in the diagram. This frequency transition is even more sharp at intermediate values of *W* (i.e. *W* = ∼20): dark blue abruptly changes to yellow. Our simulations have shown that in the areas in which frequency does not change much, the increasing *G* leads predominantly to synchronization of bursting neurons with *I*_NaP_ and involving in the populational bursting some neurons without *I*_NaP_. However, at some higher values of *G*, the remaining tonically active neurons depolarize neurons with *I*_NaP_ that have not been initially involved in bursting, leading to abrupt transitions to a higher frequency. Then again increasing *G* results predominantly in synchronization of bursting neurons, this time at a higher frequency of populational bursting. Depending on particular distribution of neuronal parameters these transitions might happen up to three times when *W* is fixed and *G* is increasing. After all neurons are involved in populational bursting and synchronized, further increase of *G* does not produce increases in frequency of populational bursting. On the contrary, at higher *G*, the length of the populational bursts is increasing and the interburst interval is decreasing, leading to smooth transition to the slower populational bursting (shown by smooth transition from the cyan/yellow to blue in **Figure 8A**) and eventually the population switches to sustained tonic activity (shown for higher *W* values by black in **Figures 8A,B**).

**Figure 8.**
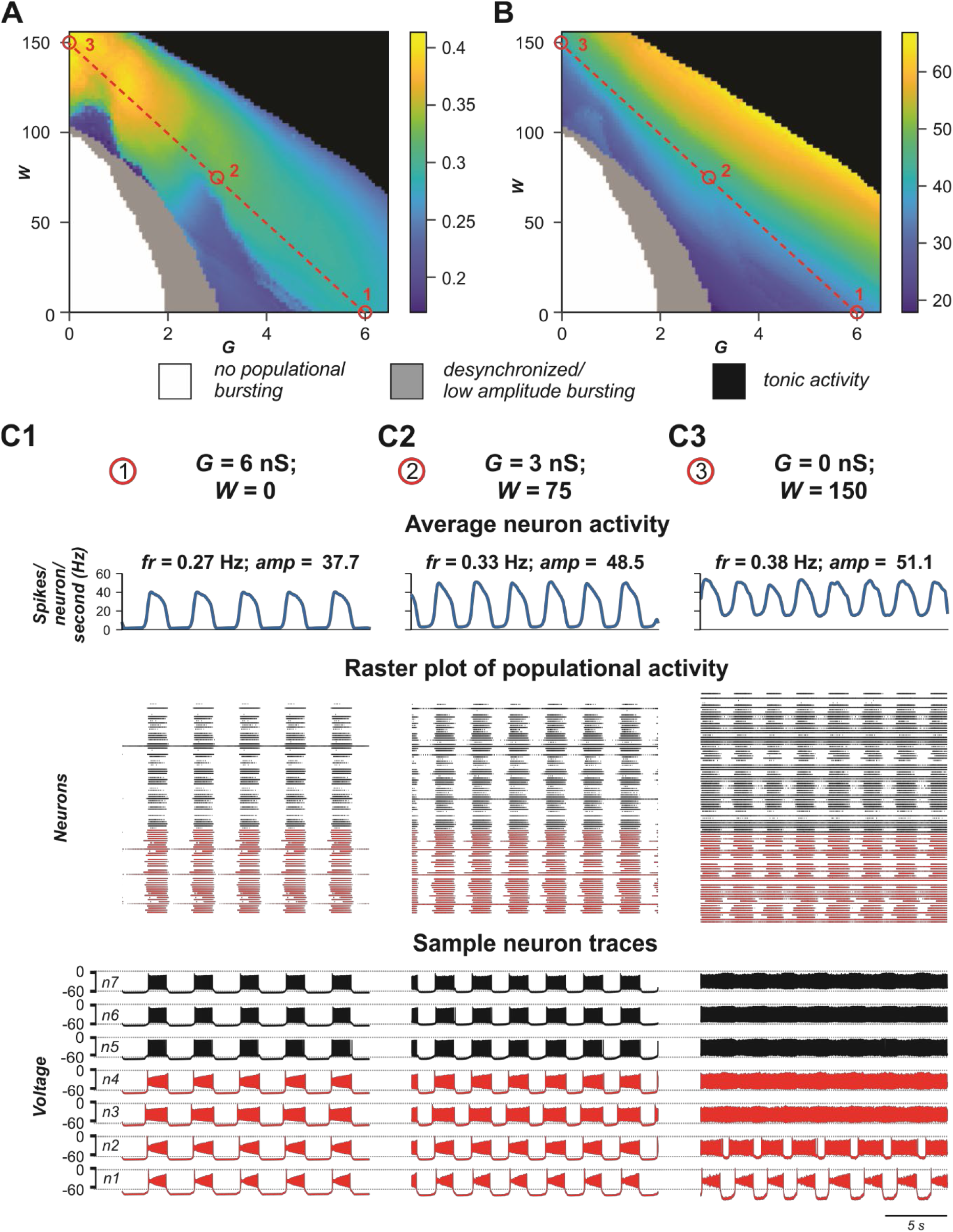
Role of electrical vs chemical coupling in generation of synchronized populational activity. **(A,B)** 2D diagrams showing color-coded frequency **(A)** and amplitude **(B)** of populational bursting when the averaged total electrical (*G*) or chemical (*W*) synaptic input to each neuron in the population was varied. **(C1-C)** Examples of simulations for three pairs of *G* and *W* values indicated in **A** and **B** by small red circles. Arrangement and notation in **C1-C3** are the same as in **Figure 4**.

With the increase of *W*, the unilateral excitatory chemical connections allow for a higher bursting frequency. If *W* is high enough, (if *W* > 20 in our simulations) the model is capable of generating synchronous populational bursting at low values of *G* or even if electrical coupling in the population is absent. With a fixed *G*, the frequency of populational bursting is increasing with an increase of the total strength of the excitatory chemical connections (*W*). However, the range of *G* in which the model generates populational bursting activity decreases with increasing *W* due to the earlier transition to sustained tonic activity of the majority neurons (black area in **Figures 8A,B**). **Figure 8B** shows that the average changes in burst amplitude depending on the values *G* and *W* are quite smooth because the amplitude depends in a greater extent on how well neuron activity in the population is synchronized and neuronal synchronization directly depends on increasing total strengths of the electrical and/or excitatory chemical connections (see also **Figures 4,7**).

**Figures 8C1-C3** demonstrate the results of simulations for three pairs of *G* and *W* values shown by small red circles in **Figures 8A,B**. In **Figure 8C1**, *W* = 0 and *G* = 6 nS. In this case, *G* is high enough to generate synchronized populational bursting activity in the absence of excitatory chemical connections. **In Figure 8C2**, *G* = 3 nS and at that strength of electrical coupling the model cannot generate synchronized populational bursting in the absence of excitatory chemical connections. However, populational bursting can be re-established with increasing *W*, which is illustrated in **Figure 8C2** for *W* = 15. On the other hand, if *W* is high enough, populational bursting activity emerges even without electrical coupling of neurons in the population as can be seen in **Figure 8C3** at *W* = 30 and *G* = 0 nS. It should be noted that in the latter case, a higher level of background activity is present.

Altogether, our simulations indicate that both electrical and excitatory chemical connections between neurons in the heterogeneous population incorporating a subset of intrinsically bursting neurons support synchronization of activities of individual neurons resulting in the rhythmic populational bursting activity. However, the effects of increasing strength of electrical and excitatory chemical coupling of frequency of populational bursting are different.

## 3 Discussion

This study is based on a previous, experimental investigation by Ha and Dougherty (2018) that revealed the presence of electrical coupling between Shox2 nonV2a interneurons, which have been suggested to be critically involved in neuronal synchronization and generation of populational bursting in the neonatal rodent spinal cords. Here, we used computational modeling to investigate the potential roles of different neural interactions (gap junctional coupling and/or excitatory chemical synapses) and their interactions with the cellular *I*_NaP_-dependent bursting properties in the neuronal synchronization and generation of locomotor-like oscillations. We simulated and analyzed the behavior of pair of neurons operating in different regimes as well as heterogeneous populations of 100 neurons in different states, sparsely connected by either gap junctions or excitatory chemical synapses, and studied the possible role of electrical and chemical synaptic connection in populational bursting and control of frequency and amplitude of populational oscillations. The different aspects of our simulation results in connection with previous experimental findings as well as limitations of current studies are discussed below.

### 3.1 The role of electrical coupling between spinal interneurons in rhythmic activity in the isolated spinal cord of neonatal rodents

Electrical coupling between neurons is known to be widely present in different regions of the central nervous system of mammals, including rodents, especially in the early stages of the development. The role of gap junctions in neuronal synchronizations and generation of populational oscillations has been previously studied using computational neuron models of different complexities (Kepler et al., 1990; Sherman and Rinzel, 1992; Pfeuty et al., 2003; Kopell and Ermentrout, 2004). Gap junctional coupling has been demonstrated to interconnect spinal Shox2 nonV2a neurons (Ha and Dougherty, 2018) as well as Hb9 neurons (via other unidentified neurons, Wilson et al., 2007). It has been suggested that electrical coupling within each of these genetically identified populations significantly contributes to neuronal synchronization and populational bursting during spinal circuit development. Although the role of this coupling in the generation of locomotor-like oscillations in neonatal mouse cords has not been directly demonstrated for the specific spinal neuronal populations, pharmacological suppression of gap junctions in the cord in general resulted in a decrease of frequency and amplitude of locomotor-like oscillations recorded from ventral roots of the neonatal spinal cord during drug-evoked fictive locomotion (Ha and Dougherty, 2018). Our simulations of a heterogeneous population of neurons connected with gap junctions have demonstrated the dependence of frequency and amplitude of populational bursting on the strength of electrical coupling and the reduction of both characteristics with the reduction of this coupling, similar to the experimental results of pharmacological suppression of gap junctions. This provides additional support for the role of electrical coupling in the locomotor activity in the spinal cords of neonatal rodents.

### 3.2 The potential role of persistent sodium current in locomotor-like oscillations in the cord and its interaction with gap junction neuronal coupling

Despite many years of intensive investigations, the role of intrinsic biophysical neuronal properties in the generation of locomotor-like oscillations in the mammalian spinal cords remains largely unknown. Based on the previous data and computational models on the generation of respiratory oscillations in the mammalian brainstem (Butera et al., 1999a,b; Del Negro et al., 2001; Rybak et al., 2004), a series of computational models of spinal locomotor circuits has been developed by suggesting the critical role of *I*_NaP_ in the generation of locomotor rhythmic activity in the spinal cord (Rybak et al., 2006a,b; McCrea and Rybak, 2007). This suggestion has been partly supported by several independent studies confirming the presence of *I*_NaP_ in the neonatal rodent spinal cords (Zhong et al., 2007; Tazerart et al., 2007, 2008; Ziskind-Conhaim et al., 2008; Harris-Warrick, 2010; Brocard et al., 2010, 2013; Tong et al., 2012). Consequently, most of the recent computational models of spinal locomotor circuits which incorporated *I*_NaP_ considered this current to be critically involved in the spinal cord rhythmogenesis (Sherwood et al., 2011; Zhong et al., 2012; Brocard et al., 2013; Rybak et al., 2013, 2015; Shevtsova et al. 2014; Danner et al., 2016, 2017, 2019; Shevtsova and Rybak, 2016; Ausborn et al., 2019). The persistent sodium channels, including their voltage-dependent activation, have been characterized in spinal Hb9 interneurons present in the spinal cord (Brocard et al., 2013) and presumably contribute to the generation and/or propagation of locomotor-like oscillations (Wilson et al., 2007; Ziskind-Conhaim et al., 2008; Brocard et al., 2013). Although *I*_NaP_ has not yet been characterized in Shox2 neurons, there is evidence for the expression of persistent inward currents in these neurons which likely includes *I*_NaP_ (Ha et al., 2019). In this study, we followed the previous computational models of the spinal locomotor circuits by suggesting that *I*_NaP_ plays critical role in the generation of locomotor-like oscillations. We, however, suggested that this current is present in, and randomly distributed over, only a subset of all neurons involved in generation of populational oscillations (with basic ratio of 40/60%). Our particular interest was in considering the behavior of neurons with and without *I*_NaP_ connected by gap junctions and the possible roles of this current and electrical coupling in initiation of synchronized bursting activity. We have shown that electrical coupling leads to equilibration of membrane potentials between the coupled neurons. Depending on the membrane potentials of neurons, the electrical coupling can increase and decrease bursting frequency and bring neurons without *I*_NaP_ from silent or tonic to bursting mode, hence amplifying neuronal synchronization. We have also shown that the presence of *I*_NaP_ in coupled neurons can amplify the effect of electrical coupling, leading to emergence of bursting activity in initially silent neurons (see **Figure 2**). This corresponds to conclusion made from other studies that *I*_NaP_ is a strong modulator of electrical synapses (Haas and Landisman, 2012). Finally, we have shown that the presence of *I*_NaP_ in some neurons in cooperation with electrical and/or chemical coupling between neurons promotes neuronal synchronization in heterogenous neuronal population and that this current can be critical for initiation and support of populational bursting. Since incorporation of *I*_NaP_ into only a subset of neurons in the modelled population was enough to generate a synchronized populational bursting when bidirectional connections at incidences seen electrophysiologically were incorporated, we would predict that to generate rhythmic locomotor-like oscillations in the neonatal spinal cord, it is sufficient for *I*_NaP_ to be present in a subset of rhythm generating neurons but not necessarily in all of them.

### 3.3 Gap junctions vs. chemical synaptic connections

Electrical transmission has been shown to be prevalent during early postnatal period but decreases as the animal matures (Walton and Navarret, 1991; Chang et al., 1999). Although the decline is seen within the first postnatal week in motor neurons (Walton and Navarret, 1991; Chang et al., 1999), gap junctional coupling within both Shox2 non-V2a neuron and Hb9 neuron populations remains relatively high throughout the first two postnatal weeks (Ha and Dougherty, 2018; Hinckley and Ziskind-Conhaim, 2006). The probability of coupling among neighboring neurons was found to be ∼0.3 (Ha and Dougherty, 2018) and this is the value that was chosen for the modeling study. The probability value of 0.1 was set for chemical synapses and falls more in line with a prior Shox2 connectivity study (Dougherty et al., 2013). There were a few key differences between the reports of Shox2 connectivity. First, the experiments showing connectivity by chemical synapses at a rate of ∼10% (Dougherty et al., 2013) was performed in mice where the Chx10 absence/presence was not distinguished, so it included the entire Shox2 population as a single sampling group. Connections were tested in a dorsal horn removed preparation and were typically within the field of view of a 60x objective but not necessarily neighboring neurons and there were no instances of electrical connections reported. In Ha and Dougherty (2018), pairs of neurons recorded were in close proximity (∼50μm on average) and were selected based on the visualization of processes crossing and in close apposition. Further, these neurons were distinguished based on the presence/absence of Chx10 expression. Therefore, the 33% incidence of electrical synapses is only among very close neighboring neurons of the same group. In this study, chemical connections were only found at 2%, indicating that neighboring neurons are less likely to be connected by chemical synapses. Both values of chemical connections are likely to be underestimated, particularly the one in which only near neighboring neurons were recorded. Interestingly, the discrepancies in the results of these studies reinforces the idea that electrical connectivity is mechanism for local synchronization.

Gap junctions behave as low pass filters and efficiently convey membrane potential fluctuations. Additionally, action potentials in one Shox2 neuron reliably lead to spikelets in gap junctionally connected neurons. However, chemical synapses between neonatal Shox2 neurons had high failure rates (Ha and Dougherty, 2018). Therefore, it is possible that there is a higher incidence of chemical synaptic connections in older ages, not due to new synapses but increased efficacy at synapses established early in development. The experimental study considered single pairs using dual recordings and responses to current injection in the presynaptic neuron. It is unlikely that the postsynaptic cell receives input from a single presynaptic Shox2 neuron. Gap junctional coupling between multiple presynaptic neurons will synchronize their firing, as demonstrated in the current study, likely leading to a more reliable excitation of the postsynaptic neuron, even if excitatory chemical synapses are sparse and single synapses have high synaptic failure rates. This could provide a mechanism for increased efficacy of excitatory transmission not only within the population but in transmission to downstream targets.

Using our model, we investigated how the activity of the population might change with changing electrical and chemical synaptic connections. Our simulations demonstrated that both electrical and excitatory chemical connections between neurons in the heterogeneous population support synchronization of activities of individual neurons resulting in the rhythmic populational bursting activity. This implicitly confirms that the locomotor-like oscillations in spinal cord can be maintained over a continuum. Thus, the results suggest that gap junctional coupling strongly drives synchronization of the population. As bidirectional electrical coupling declines with age, other mechanisms can take over to maintain rhythmic bursting within the population. The change to a dominance of unidirectional chemical synaptic interactions, particularly with *I*_NaP_ present in a subset of these neurons, is sufficient to fulfill this role and to expand the range of attainable frequencies. Interestingly, our simulation results have demonstrated that a strong, synchronous population bursting can be produced in a population when as low as 10% of neurons are connected by excitatory chemical synapses.

Our simulations have also demonstrated that the effects of increasing the strength of electrical and excitatory chemical coupling on the frequency of populational bursting are different. In both cases, a subset of neurons in the population generates spiking activity and provides an excitatory tonic input to other neurons which is enhanced with increasing connection strength and results in an increased frequency of populational bursting. In addition, in the case of bidirectional electrical coupling, tonically active neurons directly depolarize other neurons, elevating neuronal resting potentials and leading to an increase in frequency of populational bursting. However, in the case of bidirectional electrical coupling, bursting neurons have a hyperpolarizing effect on tonically firing neurons and eventually an increase of *g*_Gap_ turns spiking activity of majority of tonically firing neurons into bursting, as was shown in our simulations (see **Figures 1A3,B3** and **4**) and in earlier studies (Sherman and Rinzel, 1992). This prevents further increase in bursting frequency. After all of the neurons become involved in populational bursting and their membrane potentials reach equilibrium, further increase of *g*_Gap_ does not increase the frequency of populational oscillations (see also Kepler et al., 1990). Instead, the frequency starts to decrease. In contrast, in populations with chemical coupling, elevation of *w*_Syn_ increases the activity of tonically active neurons which depolarize bursting neurons, hence leading to the increase the frequency of populational bursting.

### 3.4 Limitations of the present studies and future directions

In this modeling study we limited single neuron models by incorporating only minimal number of ionic channels necessary for generation of neuronal spiking activity, such as fast sodium (Na) and potassium rectifier (K), and persistent sodium (NaP) involved in bursting rhythmic activity. Therefore, these models remain largely generic. At the same time, we know that spinal neurons, and particularly Shox2 neuron types contain other ionic channels, including T-type calcium and h currents and different Ca^2+^ and Ca^2+^-dependent potassium currents that can also support pacemaker properties and have been implicated in locomotor-like activity (Wilson et al., 2005; Anderson et al., 2012; Kiehn, 2016; Brocard, 2019). In addition, there is indirect evidence of an important role of Na^+^/K^+^ pump in CPG operation (Kueh et al., 2016) and particularly in the neonatal spinal cords of mice (Picton et al., 2017). The role of Na^+^/K^+^ pump current has been previously included in the computational models of Hb9 neurons (Brocard et al., 2013) and can be included in Shox2 neuron models in the future. This current is known to interact with h-current (Kuen et al., 2016) and can specifically affect neuronal coupling via electrical and chemical synapses. This awaits additional experimental and computational investigations. A more realistic model of different types of spinal interneurons that includes different ionic channels with experimentally measured characteristics should be developed and investigated (see for example, Sharples et al., 2020), which represents one of the directions of our future investigations.

The other limitation of the present study is the consideration of only excitatory interactions within the spinal network. The potential role of electrical coupling between inhibitory interneurons and the role of inhibitory network interactions in cooperation with gap junction (Bennett, 1997; Bennett and Zukin, 2004; Connors, 2017; Kopell and Ermentrout, 2004) will also be a focus of our future investigations.

The contribution of gap junctional coupling within specific populations to locomotor rhythmicity remains to be tested experimentally. The pharmacological studies are limiting in that the gap junction blockers are acting at all electrical synapses, which have been shown between several classes of interneurons (Wilson et al., 2007, Hinckley and Ziskind-Conhaim, 2006, Chopek et al., Zhong et al., 2010) and motor neurons (i.e. Walton and Navarret, 1991, Tresch and Kiehn, 2000, Rash et al., 1996, Chang et al., 1999). Additionally, carbenoxolone, the most common gap junction blocker, has been shown to have additional actions which may affect rhythmicity (Connors, 2012; Tovar et al., 2009, Elsen et al., 2008). Therefore, conditional knockout studies are required to determine implications more precisely.

Finally, the mutual interactions between different classes of genetically identified excitatory and inhibitory interneurons present in the mammalian spinal cord also awaits further experimental and computational investigations.

## 4 Materials and Methods

### 4.1 Experimental methods

*All experimental procedures followed NIH guidelines and were reviewed and approved by the Institutional Animal Care and Use Committee at Drexel University (protocols 20317 and 20657)*.

We performed further analysis of ventral root recordings during fictive locomotion (evoked with NMDA and 5-HT) prior to and after application of 100μM carbenoxolone, collected for **Figure 6** in Ha and Dougherty (2018) with detailed methods contained therein. Briefly, spinal cords were isolated neonatal mice (P0-5) in a cold dissecting solution containing in mM: 111 NaCl, 3 KCl, 11 glucose, 25 NaHCO_3_, 3.7 MgSO_4_, 1.1 KH_2_PO_4_, and 0.25 CaCl_2_, bubbled with 95%/5% O_2_/CO_2_. Cords were then transferred to a room temperature recording solution containing in mM: 111 NaCl, 3 KCl, 11 glucose, 25 NaHCO_3_, 1.3 MgSO_4_, 1.1 KH_2_PO_4_, and 2.5 CaCl_2_, bubbled with 95%/5% O_2_/CO_2_. Glass suction electrodes were used to record activity from 2-3 lumbar (L)2-5 ventral roots, band-pass filtered at 10-1,000 Hz. Locomotion was evoked by bath application of N-Methyl-D-aspartic acid (NMDA, 7 μM, Sigma) and serotonin creatinine sulfate monohydrate (5-HT, 8 μM, Sigma). The gap junction blocker carbenoxolone (Sigma) was applied at 100μM. Here, twenty-five bursts immediately before carbenoxolone application and 30 min after the start of carbenoxolone application were subjected to analysis previously described (Dougherty et al., 2013). Briefly, ventral root activity was rectified and smoothed (time constant 0.2s) in Spike2 (Cambridge Electronic Design). Recordings included either 2 ventral roots or 3 ventral roots. The cycle and amplitude values were taken from a rostral lumbar root (L1, L2, or L3) ipsilateral to a recorded caudal lumbar root (L4 or L5). The frequency was calculated from the mean cycle period, each measured from burst onset to onset. Amplitude was the measured peak amplitude of each burst. The mean values were normalized to the no drug condition to control for variations inter-experiment variation including root size, suction electrode size and seal.

### 4.2 Computational methods

#### 4.2.1 Neuron model

All neurons were simulated in the Hodgkin-Huxley style as single-compartment neuron models. The membrane potential, *V*, in neurons with *I*_NaP_ was described by the following differential equation:

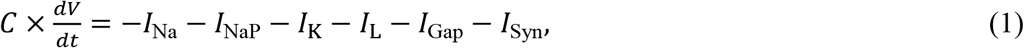

where *C* is the membrane capacitance and *t* is time.

The membrane potential, *V*, in neurons without *I*_NaP_ current was described as follows:

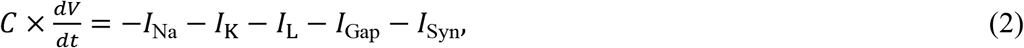

The ionic currents in Equations (1) and (2) were described as follows:

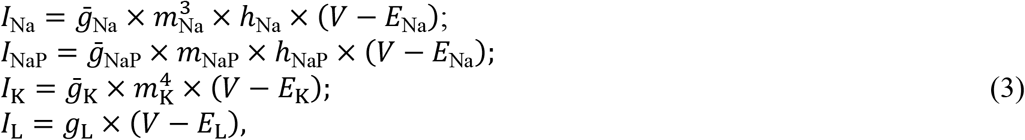

where *I*_Na_ is the fast sodium current with maximal conductance 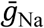; *I*_NaP_ is the persistent (slowly inactivating) sodium current with maximal conductance 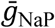; *I*_K_ is the delayed-rectifier potassium current with maximal conductance 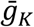; *I*_L_ is the leakage current with constant conductance *g*_*L*_. *E*_Na_, *E*_K_, and *E*_L_ are the reversal potentials for sodium, potassium and leakage currents, respectively; *m* and *h* with indexes indicating currents are, respectively, the activation and inactivation variables of the corresponding ionic channels.

Dynamics of activation and inactivation variables of voltage-dependent ionic channels (Na, NaP, and K) in Equation (3) was described by the following differential equations:

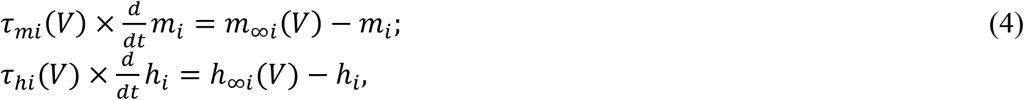

where *m*_∞*i*_(*V*) and *h*_∞*i*_(*V*) define the voltage-dependent steady-state activation and inactivation of the channel *i*, respectively, and *τ*_*mi*_(*V*) and *τ*_*hi*_(*V*) define the corresponding time constants. Activation of the fast and persistent sodium channels is was instantaneous. The expressions for channel kinetics in Equation (4) are described as follows:

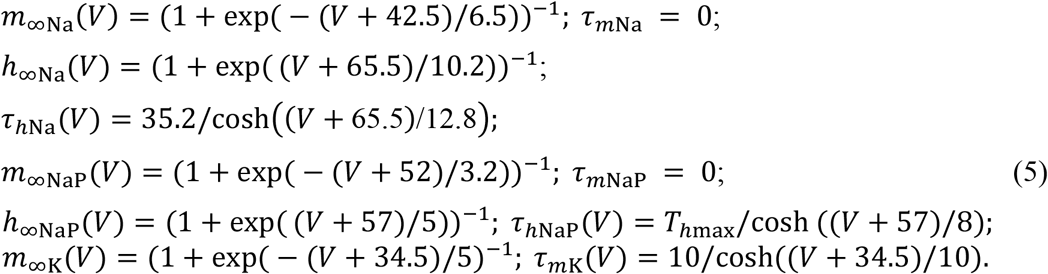

Parameters for channel kinetics were derived from Brocard et al. (2013). The maximal conductances for the fast sodium, delayed-rectifier potassium, leakage currents were, respectively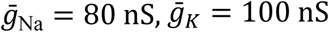, and *g*_*L*_ = 1 nS. The maximal conductance and inactivation time constant for *I*_NaP_, 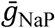 and *T*_*hmax*_, and the leakage reversal potential, *E*_L_, in different simulation were varied.

The synaptic current (*I*_Syn_ with conductance *g*_Syn_ and reversal potential *E*_Syn_) for the *j*-th neuron was described as follows:

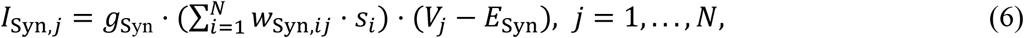

where for excitatory chemical synapses *g*_Syn_ = 1 nS, *E*_Syn_= 0, and *w*_Syn,*ij*_ = *w*_Syn_ if the *i*-th and *j*-neurons were synaptically coupled and equal to 0 otherwise. The value of *w*_Syn_ was varied.

The synaptic variable *s* was governed by first order activation scheme (Compte et al., 2005):

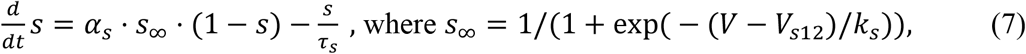

where *α*_*s*_ = 1; τ_*s*_ = 15 mS; *V*_*s*12_ = −20; and *k*_*s*_ = 2.

The electrical coupling current *I*_Gap_ for the *j*-th neuron was described as follows:

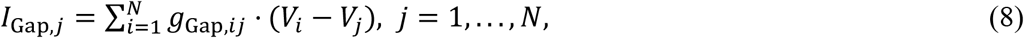

where the coupling strength *g*_Gap,*ij*_ = *g*_Gap_ if the *i*-th and *j*-th neurons share a gap junction and is equal to 0 otherwise. The value of *g*_Gap_ was varied.

The following general neuronal parameters were assigned: *C* = 40 pF; *E*_Na_ = 55 mV; *E*_K_ = − 80 mV.

#### 4.2.2 Two-cell model

Each neuron in the two-cell model represented either a bursting neuron incorporating the persistent sodium current, *I*_NaP_, or simple tonically spiking neuron without *I*_NaP_. Dynamics of the membrane potential of the neurons was described by Equations (1-5) where 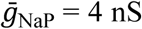, if not otherwise stated. These neurons were coupled either bidirectionally by electrical synapses as described by Equation (8) (*g*_Gap_ = {0.05, 0.1, 0.2}) or unidirectionally by AMPA synapse (see Equations (6) and (7), *w*_Syn_ = {0.5, 1, 2}). The initial operational regime of each neuron was defined by the value of its leakage reversal potential, *E*_L_, indicated in **Tables 1,2**.

#### 4.2.3 Neuron population

The neuron population contained 100 neurons and included two subpopulations: neurons incorporating *I*_NaP_ (subpopulation 1) and neurons non-incorporating *I*_NaP_ (subpopulation 2). In our simulations, *N*_NaP_, number of neurons incorporating *I*_NaP,_ varied from 10 to 40 and the observed results were qualitatively similar.

Heterogeneity of neurons within the population was provided by random distributions of the baseline values of the mean conductance of the leakage channel, *g*_*L*_, and the leakage reversal potentials, *E*_*L*0*i*_ (*i* = 1,2) for two subpopulations, mean maximal conductance of the persistent sodium channel, 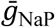, in the subpopulation of neurons incorporating *I*_NaP_, and initial conditions for the values of membrane potential and channel kinetics variables. The baseline values of distributed parameters and initial conditions were assigned prior to simulations from their defined average values and variances using a random number generator (normal distribution), and a settling period of 10-40 s was allowed in each simulation. In our simulations we varied *E*_*L*01_= {-73, -74, -75} for subpopulation of neurons with *I*_NaP_; *E*_*L*02_= {-68, -70, -72} for subpopulation of neurons without *I*_NaP_; parameter 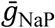 was varied in the range of [2, 5]. We also run simulation with different values of the maximal time constant of the persistent NaP ^+^ current inactivation, *T*_*hmax*_ = {7, 9, 10} seconds. In simulations for the populational model shown in this paper: *N*_NaP_ = 40; *E*_*L*01_= -74±14.8 mV; *E*_*L*02_= -70±14 mV; *g*_*L*_ = 1±0.2; 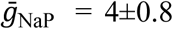; *T*_*hmax*_ = 9 s, if not stated otherwise.

Random connections between neurons in the population were assigned prior to each simulation based on probability of connections. A random number generator was used to define the existence of each synaptic connection. Typically, we used *p*_Gap_ = 0.3 for gap junctional connections and *p*_Syn_ = 0.1 for excitatory chemical connections. After the distribution of connections was set, the connection strengths of electrical and/or excitatory chemical synapses (*g*_Gap_ and *w*_Syn_) were altered. To estimate dependence of the model behavior on coupling strength and type when both types of connections were present on the model, we introduced two variables *G* and *W*, (*G* = *N* x *p*_Gap_ x *g*_*GaP*_ and *W* = *N* x *p*_Syn_ x *w*_*Syn*_), characterizing the average total network electrical or chemical synaptic input to each neuron in the population. To define electrical and chemical synapses in the populational model we used standard definition of these synapses by Brain2 (Stimberg et al., 2014, 2019). The coupling strengths for electrical and chemical connection (*g*_Gap_ and *w*_Syn_ or *G* and *W*) are specified for each simulation.

#### 4.2.4 Computer simulations

The simulations for the two-cell model were run by using a custom Matlab script (The Mathworks, Inc., Matlab 2020a). The simulations for the populational model were performed using the custom script written in Python and implemented in Brian2 environment (Stimberg et al., 2014, 2019). Differential equations were solved using the second order Runga-Kutta integration method. Simulation results were saved as the ASCI file containing the values of *g*_Gap_ and *w*_Syn_ and calculated mean values and standard deviation over simulation time for the period and amplitude of populational bursting. We also saved the figures showing populational activity, raster plot of neuron spiking, and voltage traces of sample neurons over simulation time to visually estimate the results of simulations. The scripts and model configuration files to create the simulations presented in the paper are available at the [NAME OF REPOSITORY and LINK to be added upon acceptance of manuscript].

#### 4.2.5 Data analysis in computer simulations

The results of simulations were processed by custom Matlab scripts (The Mathworks, Inc., Matlab 2020a). For each neuron period of bursting was estimated as the difference between two consecutive burst onsets and averaged for the time course of simulation. Frequency was calculated as reciprocal to the period. To assess the populational model behavior, the averaged integrated activity of neurons in the population (average number of spikes per neuron per second) was used to calculate the oscillation period. The timing of burst onsets was determined at a threshold level equal to 30% of the average populational burst amplitude in the current simulation. For some simulation with less stable populational activity we determined the threshold level by visual inspection of the results. The bursting period was defined as the duration between two consecutive burst onsets. Duration of individual simulations depended on the values of parameters *g*_Gap_ and *w*_Syn_, and to robustly estimate average value of the oscillation period, the first 10–20 transitional cycles were omitted to allow stabilization of model variables, and the values of the bursting period and amplitude were averaged for the next 10–20 consecutive cycles. To build 2D diagrams we estimated stability of the populational bursting activity by calculating values of the period and amplitude of populational bursting and their standard deviations. All simulations were divided in four groups: no populational activity, low amplitude (less than 10 spikes per neuron per second) or highly unstable populational activity (SD of calculated period ≥ 50%), stable populational bursting, and tonic populational activity. For the stable populational bursting, frequency of populational oscillations was determined as reciprocal to the mean period. The calculated frequency and amplitude were color-coded to build 2D diagrams in G x *W* parameter space. Re-initialization of the randomized parameters within the same ranges resulted in qualitatively similar simulation results.

## 6 Author Contributions

NAS, NTH, IAR, and KJD: conceptualization and investigation. NAS, IAR, and KJD: writing (original draft). NAS, NTH, IAR, and KJD: writing (review and editing). NTH and KJD: experiments and data curation. NAS: modeling, formal analysis and software. NAS and KJD: visualization. IAR and KJD: supervision, project administration, and funding acquisition.

## 7 Acknowledgments

This study was supported by grants from the National Institutes of Health (R01 NS090919; R01 NS095366; R01 NS100928; R01 NS110550). We are grateful to Simon Danner and Lihua Yao for technical assistance.

This manuscript has been released as a pre-print at https://www.biorxiv.org/content/10.1101/2020.09.15.298281v1 (Shevtsova et al., 2020).

## 8 Data Availability Statement

The experimental and modeling datasets for this study can be found in the [NAME OF REPOSITORY] [LINK] [will be uploaded after revision].

